# Mesolimbic dopamine D2 receptors and neural representations of subjective value

**DOI:** 10.1101/718858

**Authors:** Jaime J. Castrellon, Jacob S. Young, Linh C. Dang, Ronald L. Cowan, David H. Zald, Gregory R. Samanez-Larkin

## Abstract

The process by which the value of delayed rewards is discounted varies from person to person. It has been suggested that these individual differences in subjective valuation of delayed rewards are supported by mesolimbic dopamine D2-like receptors (D2Rs) in the ventral striatum. However, no study to date has documented an association between direct measures of dopamine receptors and neural representations of subjective value in humans. Here, we examined whether individual differences in D2R availability were related to neural subjective value signals during decision making. Human participants completed a monetary delay discounting task during an fMRI scan and on a separate visit completed a PET scan with the high affinity D2R tracer [18F]fallypride. Region-of-interest analyses revealed that D2R availability in the ventral striatum was positively correlated with subjective value-related activity in the ventromedial prefrontal cortex and midbrain but not with choice behavior. Whole-brain analyses revealed a positive correlation between ventral striatum D2R availability and subjective value-related activity in the left inferior frontal gyrus. These findings are the first to identify a link between directly-measured mesolimbic dopamine function and subjective value representation in humans and suggest a mechanism by which individuals vary in neural representation of discounted subjective value.

## Introduction

Nearly all behavioral decisions involve judgements about the value of desired outcomes. Intuitively, all animals should choose actions that maximize outcome values when comparing multiple options that vary in costs and benefits. However, animals, including humans, vary in their decision preferences. For example, to some individuals, the subjective value of a small, certain outcome exceeds the subjective value of a much larger, uncertain outcome even if the expected value (i.e., probability of obtaining a reward multiplied by the reward amount) of the uncertain option is numerically greater. Similarly, humans regularly spend money now that would have much more spending power later if saved and invested. This tendency to discount the future such that the subjective value of a larger, delayed reward is lower than a smaller reward available now is common across many animal species.

Neuroimaging research has shown that although similar networks of regions represent subjective value across individuals, both behavioral preferences and neural representations of subjective value are also highly variable between people^1^. What, then, accounts for differences between people? Some have suggested that specific neurotransmitters, such as dopamine (DA), may influence subjective value computation and account for variation in neural representations^1–3^. While many studies using non-human primates or rodents have linked direct measurement of DA levels or the activity of DA-releasing cells to discounting behavior and subjective value coding^4^, no study to date has explored individual differences in direct measures of dopamine function and neural representations of subjective value.

Functional MRI (fMRI) studies have consistently shown that subjective value is reflected in modulation of brain activation in a network of regions including the ventromedial prefrontal cortex (vmPFC), ventral striatum (VS), and posterior cingulate cortex (PCC)^5,6^. Since subjective value scales with DA signals in the VS in nonhuman models, DA measures might vary with individual differences in subjective valuation. Direct recordings from midbrain DA neurons in monkeys and rodents provide evidence that DA neurons are sensitive to the subjective value of rewards over decreasing delays^7–9^. Providing indirect support for this mesolimbic DA narrative in humans, pharmacological manipulation of D2Rs in humans impacts delay discounting behavior^10^. However, the regional non-specificity of drugs that target D2Rs^11^ limits attempts to detail the role of specific regions in this circuit, especially since D2Rs are present across the striatum and cortex^12,13^.

Although these studies demonstrate the impact of DA on discounting behavior, less is known about how it impacts neural subjective value signals. There are two signaling pathways that may account for effects of DA on discounting: (1.) the corticostriatal loop^14^ may account for mesolimbic DA influences on prefrontal inputs to the VS^15,16^ and (2.) the ventral striatopallidal loop may account for interactions between the VS and midbrain DA^17,18^. Potentially, individual differences in the components of either of these signaling pathways might underlie differences in prefrontal, striatal, or midbrain value computation. We therefore hypothesized that individual differences in D2R availability would correlate with subjective value signals in regions encoding subjective value. Based on previous research with both human and non-human animals, we had the strongest predictions for associations of D2Rs in the ventral striatum and midbrain with subjective value signals in ventral striatum, midbrain, and vmPFC. However, we explored multiple potential associations between the VS, midbrain, vmPFC, and PCC due to evidence from functional neuroimaging studies for subjective value signals in the PCC as well^5^.

In this study, healthy young adults completed a delay discounting task for monetary rewards during an fMRI scan. On a separate visit, we collected direct measures of DA D2R availability using positron emission tomography (PET) combined with the high affinity D2R ligand [18F]fallypride. We examined whether individual differences in measures of D2R availability were related to discounted subjective value representations in the brain.

## Methods

### Participants and screening procedures

Twenty-five healthy young adults (ages 18-24, M=20.9, SD=1.83, 13 females) were recruited from Vanderbilt University, Nashville, TN in 2012. Participants were subject to the following exclusion criteria: any history of psychiatric illness on a screening interview (a Structural Interview for Clinical DSM-IV Diagnosis was available for all subjects and confirmed no history of major Axis I disorders)^19^, any history of head trauma, any significant medical condition, or any condition that would interfere with MRI (e.g., inability to fit in the scanner, claustrophobia, cochlear implant, metal fragments in eyes, cardiac pacemaker, neural stimulator, and metallic body inclusions or other contraindicated metal implanted in the body). Participants with major medical disorders including diabetes and/or abnormalities on screening comprehensive metabolic panel or complete blood count were excluded. Participants were also excluded if they reported a history of substance abuse, current tobacco use, alcohol consumption greater than 8 ounces of whiskey or equivalent per week, use of psychostimulants (excluding caffeine) more than twice at any time in their life or at all in the past 6 months, or any psychotropic medication in the last 6 months other than occasional use of benzodiazepines for sleep. Any illicit drug use in the last 2 months was grounds for exclusion, even in participants who did not otherwise meet criteria for substance abuse. Urine drug screens were administered, and subjects testing positive for the presence of amphetamines, cocaine, marijuana, PCP, opiates, benzodiazepines, or barbiturates were excluded. Female participants had negative pregnancy tests both at intake and on the day of the PET scan. Discounting behavioral measures for all participants in this sample were previously reported as a subsample of multiple data sets^2^ and the present analysis is comprised of data from participants with valid PET and fMRI data.

Approval for the [18F]fallypride study protocol was obtained from the Vanderbilt University Human Research Protection Program and the Radioactive Drug Research Committee. All participants completed written informed consent and study procedures were approved by the Institutional Review Board at Vanderbilt University in accordance with the Declaration of Helsinki’s guidelines for the ethical treatment of human participants.

### Delay discounting task

The delay discounting task was adapted from a previously used paradigm^20^. On each trial, participants chose between an early reward and a late reward. The delay of the early reward was set to today, 2 weeks, or 1 month, while the delay of the late reward was set to 2 weeks, 1 month, or 6 weeks later. The early reward magnitude ranged between 1% and 50% less than the late reward (determined by a Gaussian distribution, min=$5, max=$30, mean=$15, standard deviation=$10). Participants played 84 trials (42 trials in two runs) of the task. Participants had up to 8 seconds to select a reward, after which their selection was highlighted for 2 seconds, followed by an inter-trial-interval that lasted up to 10 seconds minus the reaction time. If no response was made on the choice slide within 7950 milliseconds, the blank (ITI) slide was set to 2050 milliseconds. Thus, each trial lasted approximately 12 seconds from the choice screen onset to the end of an ITI. (See Figure 1A). To ensure participants were motivated in their choices, the task was incentive compatible. Participants were instructed to treat all decisions as real because a random trial would be selected for actual payout at the end of the experiment. Participants were paid in Amazon.com credit that was emailed to them either that afternoon (if the participant selected a reward available today) or scheduled to be delivered to them at the delayed date (if the participant selected a reward available later).

**Figure 1.**
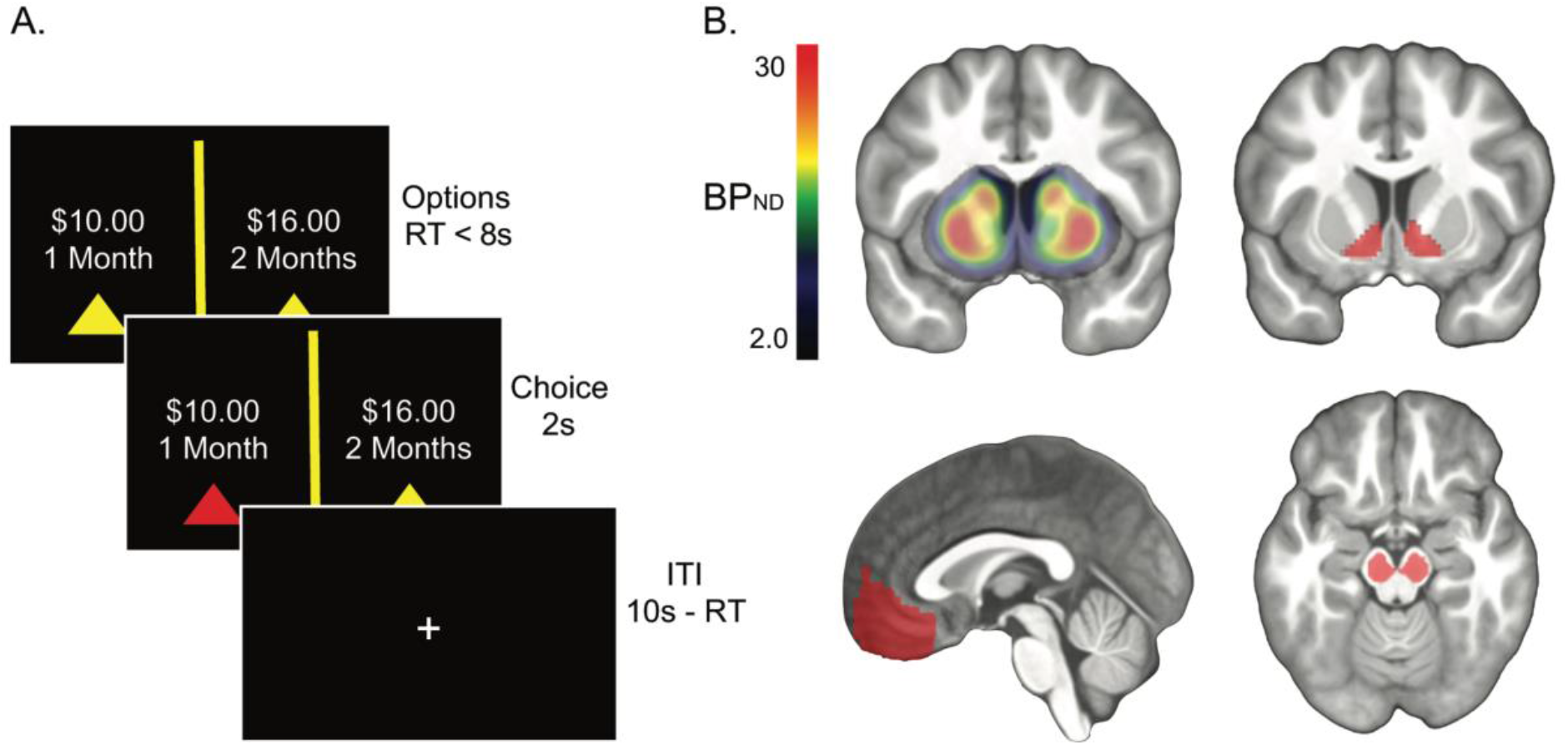
**A.)** Task outline. Participants were given up to 8 seconds to indicate a preference for a smaller-sooner or a larger-later monetary reward, after which their choice was highlighted for two seconds. Choice trials were separated by an inter-trial-interval (ITI) scaled by the difference between 10 seconds and the choice response time so that every trial lasted 12 seconds from choice onset to ITI. **B.) Top Row:** Mean BP_ND_ map (left) and ventral striatum ROI (right) from which the average DA D2 receptor availability was extracted for each subject. **Bottom Row:** Ventromedial prefrontal cortex (left) and midbrain (right) ROIs from which subjective value parameter estimates and DA D2 receptor availability were extracted for each participant. BP_ND_ map and ROIs shown are overlaid on the mean participant T1-weighted image in MNI space.

### Subjective value modeling

We used a computational model to estimate subjective value from behavioral preferences to create timeseries regressors for fMRI analysis. Since we assumed preferences were discounted hyperbolically^21,22^, for each participant, discounting was modeled with a hyperbolic discounted value function SV = A/(1+*k*D), where A represents the monetary reward magnitude, *k* represents the discount rate, D represents the delay in days, and SV represents the subjective value of available options. Choices were fit using a decision function with a softmax slope to describe how differences in subjective values between options influenced the probability of choosing the higher subjective value option. Since *k* values are not normally-distributed, we used the natural log-transformed values Ln(*k*+1) for behavioral correlations.

### PET acquisition and processing

[18F]fallypride, (S)-N-[(1-allyl-2-pyrrolidinyl)methyl]-5-(3[18F]fluoropropyl)-2,3-dimethoxybenzamide, was produced in the radiochemistry laboratory attached to the PET unit at Vanderbilt University Medical Center, following synthesis and quality control procedures described in US Food and Drug Administration IND 47,245. PET data were collected on a GE Discovery STE (DSTE) PET scanner (General Electric Healthcare, Chicago, IL, USA). The scanner had an axial resolution of 4 mm and in-plane resolution of 4.5 to 5.5 mm FWHM at the center of the field of view. Serial scan acquisition was started simultaneously with a 5.0 mCi (185 MBq) slow bolus injection of the DA D2/3 tracer [18F]fallypride (median specific activity = 5.33 mCi). CT scans were collected for attenuation correction prior to each of the three emission scans, which together lasted approximately 3.5 hours with two breaks for participant comfort.

### [18F]fallypride binding potential (BP_ND_) image calculation

Voxelwise D2/D3 binding potential images were calculated using the simplified reference tissue model, which has been shown to provide stable estimates of [18F]fallypride BP_ND_^23^. The cerebellum served as the reference region because of its relative lack of D2/D3 receptors^13^. The cerebellar reference region was obtained from an atlas provided by the ANSIR laboratory at Wake Forest University. Limited PET spatial resolution introduces blurring and causes signal to spill onto neighboring regions. Because the cerebellum is located proximal to the substantia nigra and colliculus, which both have D2Rs, only the posterior 3/4 of the cerebellum was included in the region of interest (ROI) to avoid contamination of [18F]fallypride signal from the midbrain nuclei. The cerebellum ROI also excluded voxels within 5 mm of the overlying cerebral cortex to prevent contamination of cortical signals. The bilateral putamen ROI, drawn according to established guidelines^24^ on the MNI brain, served as the receptor rich region in the analysis. The cerebellum and putamen ROIs were registered to each participant’s T1-weighted anatomical image using FSL non-linear registration of the MNI template to the individual participant’s T1. T1 images and their associated cerebellum and putamen ROIs were then co-registered to the mean image of all realigned frames in the PET scan using FSL-FLIRT (http://www.fmrib.ox.ac.uk/fsl/, version 6.00). Emission images from the 3 PET scans were merged temporally into a 4D file. To correct for motion during scanning and misalignment between the 3 PET scans, all PET frames were realigned using SPM8 (www.fil.ion.ucl.ac.uk/spm/) to the frame acquired 10 minutes post injection. Model fitting and BP_ND_ calculation were performed using PMOD Biomedical Imaging Quantification software (PMOD Technologies, Switzerland). Binding potential images represent the ratio of specifically bound ligand ([18F]fallypride in this study) to its free concentration.

The bilateral midbrain and ventral striatum ROIs were drawn in MNI standard space using previously described guidelines^24–26^ and registered to PET images using the same transformations used in BP_ND_ calculation (see Figure 1B). An additional ROI for the medial frontal cortex in MNI space was derived from the Harvard-Oxford Atlas and registered to PET images using the same transformations used in BP_ND_ calculation.

### MRI data acquisition

Brain images were collected using a 3T Phillips Intera Achieva whole-body MRI scanner using a 32-channel head coil (Philips Healthcare, Best, The Netherlands). For each run of the delay discounting task, we used T2*-weighted gradient echo-planar imaging (EPI) to acquired 262 volumes of 38 ascending slices, 3.2 mm thick with .35 mm gap (in-plane resolution 3 × 3 mm), FOV = 240 mm × 240 mm, flip angle (FA) = 79, TR = 2000 ms, TE = 35 ms. A high resolution T1-weighted image (TFE SENSE protocol; 150 slices (in-plane resolution 1 × 1 mm), FOV = 256 × 256, FA = 8, TR = 8.9 ms, TE = 4.6 ms) was acquired for registration purposes and ROI definition.

### fMRI data preprocessing

Data preprocessing was performed using fMRIPrep version 1.0.0-rc9^27^, a Nipype^28^ based tool. Each T1-weighted volume was corrected for bias field using N4BiasFieldCorrection v2.1.0^29^ and skull-stripped using antsBrainExtraction.sh v2.1.0 (using OASIS template). Cortical surface was estimated using FreeSurfer v6.0.0^30^. The skull-stripped T1-weighted volume was co-registered to a skull-stripped ICBM 152 Nonlinear Asymmetrical template version 2009c^31^ using a nonlinear transformation implemented in ANTs v2.1.0^32^.

Functional data was slice time corrected using AFNI^33^ and motion corrected using MCFLIRT v5.0.9^34^. “Fieldmap-less” distortion correction was performed by co-registering the functional image to the same participant’s T1w image with its intensity inverted^35,36^ and constrained with an average fieldmap template^37^, implemented with antsRegistration (ANTs). This was followed by co-registration to the corresponding T1-weighted volume using boundary-based registration^38^ with 9 degrees of freedom, implemented in FreeSurfer v6.0.0. Motion correcting transformations, T1-weighted transformation and MNI template warp were applied in a single step using antsApplyTransformations v2.1.0 with Lanczos interpolation.

Three tissue classes were extracted from T1w images using FSL FAST v5.0.9^39^. Frame-wise displacement^40^ was calculated for each functional run using Nipype. For more details of the pipeline see https://fmriprep.readthedocs.io/en/latest/workflows.html.

We performed voxelwise nuisance signal removal using publicly-available scripts (https://github.com/arielletambini/denoiser) to clean the data. Specifically, we denoised the data for 10 fMRIPrep-derived confounds: CSF, white matter, standardized DVARS, framewise displacement (over 0.5 mm), and six motion parameters.

FSL FEAT (www.fmrib.ox.ac.uk/fsl) was run for each participant with fixed effects across runs. Functional data were high-pass filtered with a cutoff of 100 seconds, spatially-smoothed with a 5 mm full-width-at-half-maximum (FWHM) Gaussian kernel, and grand-mean intensity normalized. FSL FILM pre-whitening was carried out for autocorrelation correction. Events were convolved with a double-gamma hemodynamic response function. A general linear model was fit to the data with a regressor for the mean (un-modulated) signal over the duration of the choice period and a regressor for the parametric modulation of subjective value at the choice reaction time with a duration of zero seconds. We applied temporal filtering and added the temporal derivative to the waveform. Visual inspection of data quality using outputs from MRIQC^41^ suggested one participant had fMRI scans with strong artefactual features. This participant was excluded from analysis. One participant was excluded from analyses because this person only had data for a single run of the task. One participant was excluded for corrupted fMRI data. This provided a final sample of 22 participants. Participant demographics and characteristics are listed in Table 1.

**Table 1.** Study sample characteristics

| Variable | Mean ± SD |
| --- | --- |
| N | 22 |
| Sex | 12 F, 10 M |
| Age | 20.9 ± 1.95 |
| Race/Ethnicity | 13 White |
| Years Education | 14.7 ± 1.43 |
| Prop(sooner) chosen | .550 ± .212 |
| Ln(*k*) | -4.68 ± 1.27 |
| Inverse temperature | 3.74 ± 4.41 |
| LLH | 11.8 ± 5.90 |
| BIC | 31.1 ± 11.8 |
| Ventral Striatum BP<sub>ND</sub> | 17.3 ± 2.90 |
| Midbrain BP<sub>ND</sub> | 1.50 ± .236 |

### Statistical analyses

Linear regressions between D2R BP_ND_ in each of the 3 PET ROIs (VS, midbrain, vmPFC) and discounting behavior (indexed with Ln(*k*+*1*) or proportion of smaller-sooner choices) was run in JASP (Version 0.9.2)^42^. For associations between PET ROIs and fMRI subjective value signal, we extracted the mean subjective value parameter estimates (percent signal change) for each participant from ROIs in the medial frontal cortex, posterior cingulate, midbrain, and ventral striatum (defined using the same methods described above for the PET ROIs) (see Figure 1B). Like the medial frontal cortex ROI, the posterior cingulate was derived from the Harvard-Oxford Atlas. Statistically-significant relationships were defined using a Bonferroni-correction for 12 tests (3 PET ROIs: VS, midbrain, vmPFC by 4 fMRI ROIs: VS, midbrain, vmPFC, PCC) on an alpha of .05 (p < .004). Effects surviving correction for multiple comparisons were followed-up with regressions controlling for age and sex as covariates of no interest. All regression coefficients reported are standardized.

For D2R ROIs significantly associated with subjective value signal, we conducted whole-brain analyses of the fMRI data to better localize the effects or identify associations in other regions. All whole-brain fMRI analyses were carried out in FSL FEAT with mixed effects using FLAME 1. Statistical maps were thresholded using a cluster-forming threshold with a height of Z > 2.3, and cluster-corrected significance of *p* < .05. Analyses were run to examine: (1) the mean effect of subjective value parametric modulation of the fMRI BOLD signal across all participants and (2) the correlation between individual differences in BP_ND_ and subjective value parametric modulation of the BOLD signal.

## Results

### Dopamine D2Rs and delay discounting behavior

As expected, computationally-derived discount rates, Ln(*k*+1), were strongly positively correlated with the proportion of smaller-sooner options chosen (β = .911, 95% CI [.794, .963], *p* <.001). As already reported in a previous publication^2^, D2R BP_ND_ was not correlated with the proportion of smaller-sooner choices or Ln(*k*+1) values for any ROI: VS (prop sooner: β = .036, 95% CI [−.391, .451], *p* = .873; Ln(*k*): β = −.021, 95% CI [−.439, .404], *p* = .927), Figure 2D; midbrain (prop sooner: β = .036, 95% CI [−.392, .451], *p* = .874; Ln(*k*): β = −.054, 95% CI [−.465, .376], *p* = .810), and vmPFC (prop sooner: β = .218, 95% CI [−.224, .586], *p* = .330; Ln(*k*): β = .195, 95% CI [−.247, .570], *p* = .384). These D2R-discounting behavior results presented here are based on a subset of the data used in the prior publication that showed no significant associations between D2R and discounting behavior in healthy adults^2^.

**Figure 2.**
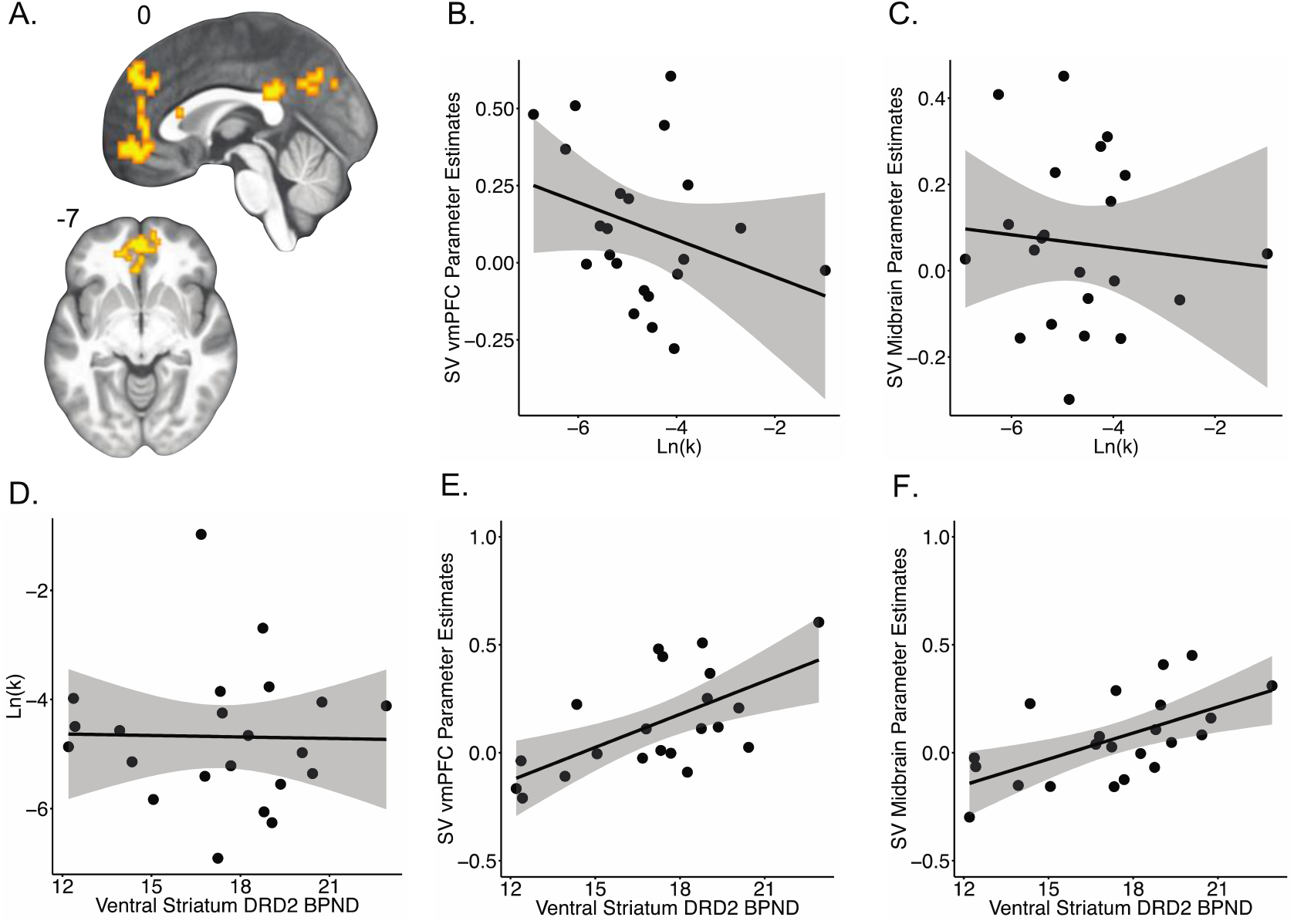
**A.)** Mean effect of subjective value (*N* = 21) overlaid on the mean participant T1-weighted image in standard space, whole brain cluster-forming threshold Z > 2.3, cluster-corrected *p* < .05. Delay discounting was not correlated with the effect of subjective value on fMRI signal in the **B.)** vmPFC (N = 22, r = −.312, *p* = .158) or **C.)** midbrain (N = 22, r = −.097, *p* = .669). DA D2-like receptor availability in the ventral striatum was not correlated with **D.)** delay discounting (N = 22, r = −.053, *p* =.821). DA D2-like receptor availability was positively correlated with the effect of subjective value on fMRI signal in the **E.)** vmPFC (N = 21, r = .624, *p* = .003) and **F.)** midbrain (N = 22, r = .597, *p* = .003). Shaded regions indicate 95% confidence interval.

### Localization of subjective value representations within fMRI data

Voxelwise analysis of the mean effect of subjective value of the chosen option revealed significant parametric modulation in the dorsomedial PFC. Exclusion of a single outlier revealed stronger and spatially extended activation in the vmPFC and posterior cingulate cortex (PCC) (see Table 2 and Figure 2A). These effects are consistent with previous studies using subjective value as a parametric regressor^6,21,43^. Both unthresholded maps with and without the outlier are available to view/download on Neurovault (https://neurovault.org/collections/PDSRXDAH/).

**Table 2.**
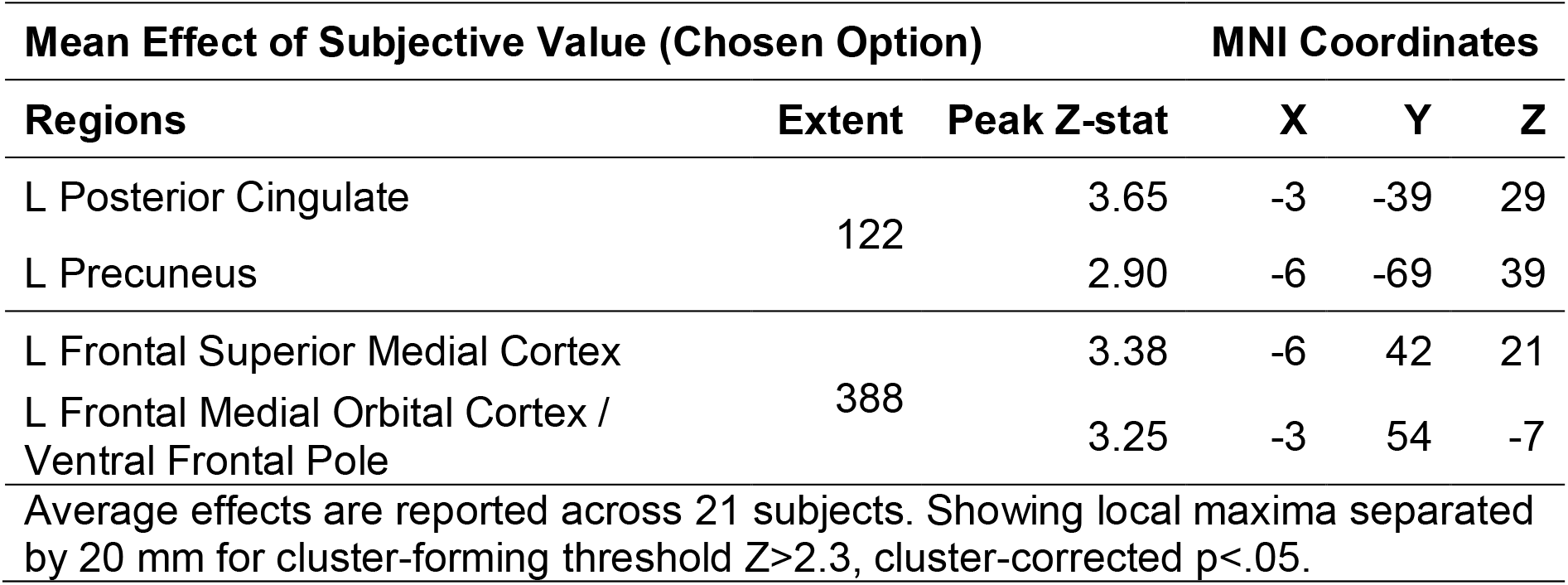
Average neural representations of subjective value of the chosen option.

### Subjective value representations and delay discounting behavior

We ran correlations between fMRI parameter estimates for subjective value in each of the 4 subjective value ROIs (VS, midbrain, vmPFC, PCC) and discounting rates to evaluate whether individual differences in subjective value representations were associated with individual differences in time discounting behavior. There were non-significant correlations between discounting and subjective value in the VS (prop sooner: β = −.046, 95% CI [−.459, .383], *p* = .839; Ln(*k*): β = −.046, 95% CI [−.383, .458], *p* = .840), midbrain (prop sooner: β = −.060, 95% CI [−.470, .371], *p* = .790; Ln(*k*): β = −.097, 95% CI [−.498, .339], *p* = .669), vmPFC (prop sooner: β = −.299, 95% CI [− .640, .140], *p* = .176; Ln(*k*): β = −.312, 95% CI [−.648, .126], *p* = .158), and PCC (prop sooner: β = −.014, 95% CI [−.433, .410], *p* = .952; Ln(*k*): β = −.009, 95% CI [−.429, .414], *p* = .967) (See Figure 2B and 2C).

### Ventral Striatum D2Rs (PET) and subjective value representations (fMRI)

We identified a positive correlation between D2R BP_ND_ in the VS and subjective value-related fMRI signal in the vmPFC (β = .466, 95% CI [.056, .742], *p* = .029). Combined visual inspection of the correlation and bivariate outlier statistics (Cook’s distance greater than 4 times the mean distance, t-test of studentized residuals (*p* < .05), and test of heteroskedasticity (*p* < .05)) identified an influential outlier with high D2R BP_ND_ but low subjective value parameter estimates that may have biased the estimated effect. This is the same outlier mentioned above in the fMRI analyses. Exclusion of this outlier revealed a stronger association (β = .624, 95% CI [.263, .832], *p* = .003) (see Figure 2E). This effect remained significant after controlling for age and sex as covariates of no interest (β = .548, *p* = .003). D2R BP_ND_ in the VS was also significantly positively associated with subjective value-related fMRI signal in the midbrain (β = .597, 95% CI [.234, .814], *p* = .003) (see Figure 2F). This effect remained significant after controlling for age and sex (β = .576, SE = .012, t(18) = 3.27, *p* = .004). D2 BP_ND_ in the VS was not significantly associated with subjective value in the VS (β = .333, 95% CI [−.103, .661], *p* = .130) or PCC (β = .026, 95% CI [−.400, .443], *p* = .909).

Exploratory voxelwise analysis of the fMRI data using ventral striatal D2R BP_ND_ revealed a significant correlation between BP_ND_ and subjective value representation in the left precentral gyrus, inferior frontal gyrus (IFG), and insula (See Figure 3, Table 3).

**Figure 3.**
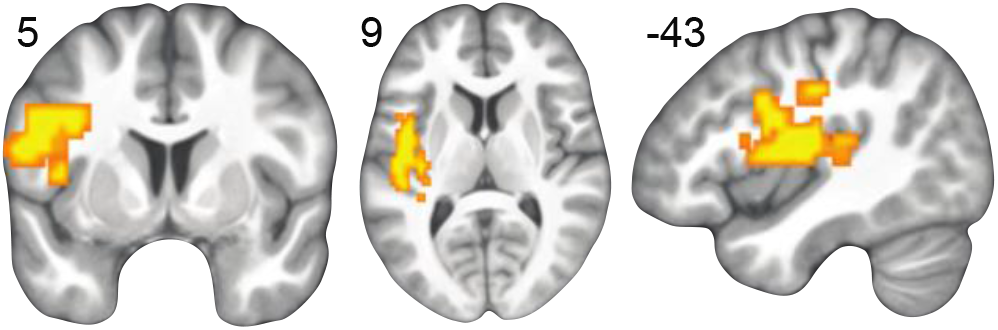
Positive correlation between ventral striatum D2 BP_ND_ and subjective value in the left inferior frontal gyrus, shown on the mean participant T1-weighted image in MNI space, whole brain cluster-forming threshold Z>2.3, cluster-corrected p<.05 shown on the mean participant T1-weighted image in MNI space.

**Table 3.**
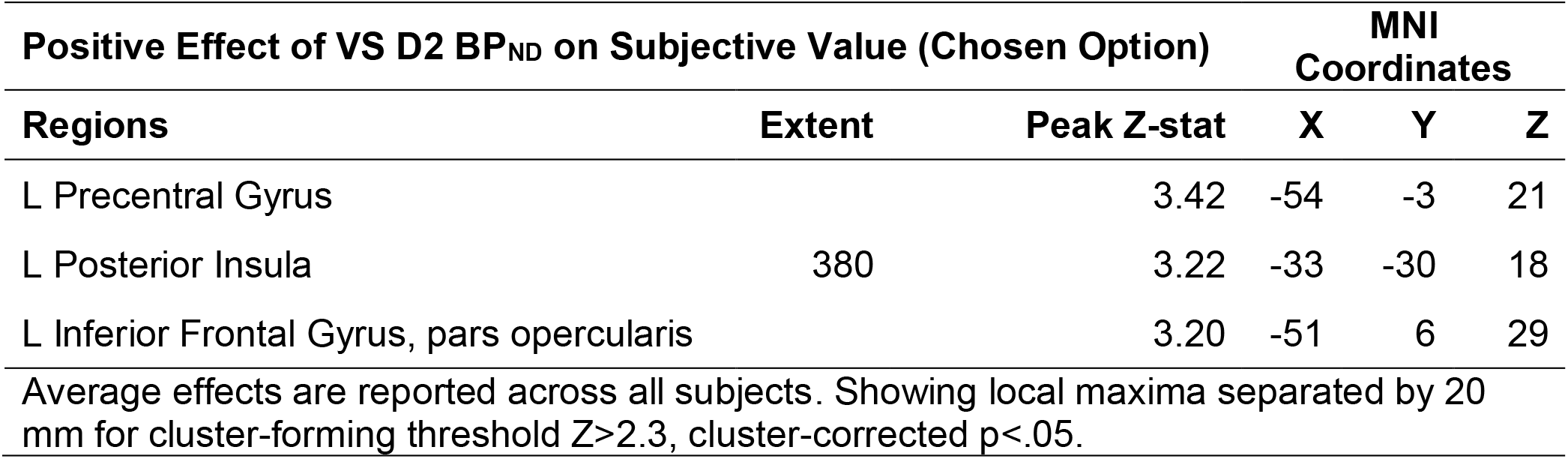
Positive correlation between ventral striatum D2 BP_ND_ and neural representations of subjective value of the chosen option.

| Positive Effect of VS D2 BP<sub>ND</sub> on Subjective Value (Chosen Option) |  |  | MNI Coordinates |  |  |
| --- | --- | --- | --- | --- | --- |
| Regions | Extent | Peak Z-stat | X | Y | Z |
| L Precentral Gyrus |  | 3.42 | -54 | -3 | 21 |
| L Posterior Insula | 380 | 3.22 | -33 | -30 | 18 |
| L Inferior Frontal Gyrus, pars opercularis |  | 3.20 | -51 | 6 | 29 |
| Average effects are reported across all subjects. Showing local maxima separated by 20 mm for cluster-forming threshold $Z > 2.3$, cluster-corrected $p < .05$. | | | | | |

### Midbrain D2Rs and subjective value representations

Midbrain D2R availability and subjective value-related fMRI signal in the vmPFC were not related (β = .254, 95% CI[−.187, .610], *p* = .254). As before, combined visual inspection of the correlation and bivariate outlier statistics (Cook’s distance greater than 4 times the mean distance, t-test of studentized residuals (*p* < .05), and test of heteroskedasticity (*p* < .05)) identified an influential outlier (same as above) with high D2R BP_ND_ but low subjective value parameter estimates that may have biased the estimated effect. Exclusion of this outlier revealed a stronger association (β = .513, 95% CI [.105, .773], *p* = .017) but the effect did not survive correction for multiple comparisons. D2 BP_ND_ in the midbrain was not significantly associated with subjective value in the midbrain (β = .398, 95% CI [−.029, .702], *p* = .067), ventral striatum (β = .026, 95% CI [−.400, .443], *p* = .909), or PCC (β = −.172, 95% CI [−.554, .269], *p* = .443). We did not conduct exploratory voxelwise analysis of the fMRI data using midbrain BP_ND_.

### Prefrontal D2Rs and subjective value representations

vmPFC D2R availability was not significantly associated with subjective value-related fMRI signal in the VS (β = −.127, 95% CI[−.521, .311], *p* = .573), midbrain (β = .143, 95% CI[−.296, .533], *p* = .525), vmPFC (β = −.129, 95% CI[−.522, .309], *p* = .567), or PCC (β = −.041, 95% CI[−.454, .388], *p* = .858). We did not conduct exploratory voxelwise analysis of the fMRI data using vmPFC BP_ND_.

## Discussion

Here we tested the hypothesis that individual differences in mesolimbic DA D2Rs relate to neural representations of subjective value. We predicted associations between D2R in ventral striatum and midbrain and subjective value signals in ventral striatum, midbrain, and vmPFC. We identified a positive correlation between VS D2R availability and the strength of subjective value signals in the vmPFC and midbrain. However, neither D2R availability nor functional neural representation of subjective value were directly correlated with discounting behavior.

The positive correlations between mesolimbic D2Rs and subjective value in the vmPFC and midbrain are consistent with past findings converging on two key circuits: a corticostriatal loop and a ventral striatopallidal loop. The corticostriatal loop is comprised of a series of pathways that promote approach behavior. Activation of D2Rs in the ventral striatum increases GABAergic signaling to the ventral pallidum which projects to the thalamus^44,45^. Neurons in the vmPFC receive these thalamic projections and promote local release of DA in the ventral striatum^46^. The ventral striatopallidal loop is comprised of connections linking the ventral striatum and dopaminergic midbrain that promote reward “wanting”^47^. Specifically, D2-mediated ventral pallidal signals from the ventral striatum that complete the corticostriatal loop also promote DA release to the ventral striatum via GABAergic signals to the midbrain^18,48^ (See Figure 4 for an illustration of these two potential mechanisms). Prevention of hyperdopaminergic states in these loops are regulated by dopamine transporters in the ventral striatum and somatodendritic autoreceptors in the midbrain^49^. Importantly, we did not measure DA release specifically in this study. Further studies with multiple measures of DA function are needed to test the specific links between subjective reward valuation and integration of value signals between the corticostriatal and ventral striatopallidal circuits.

**Figure 4.**
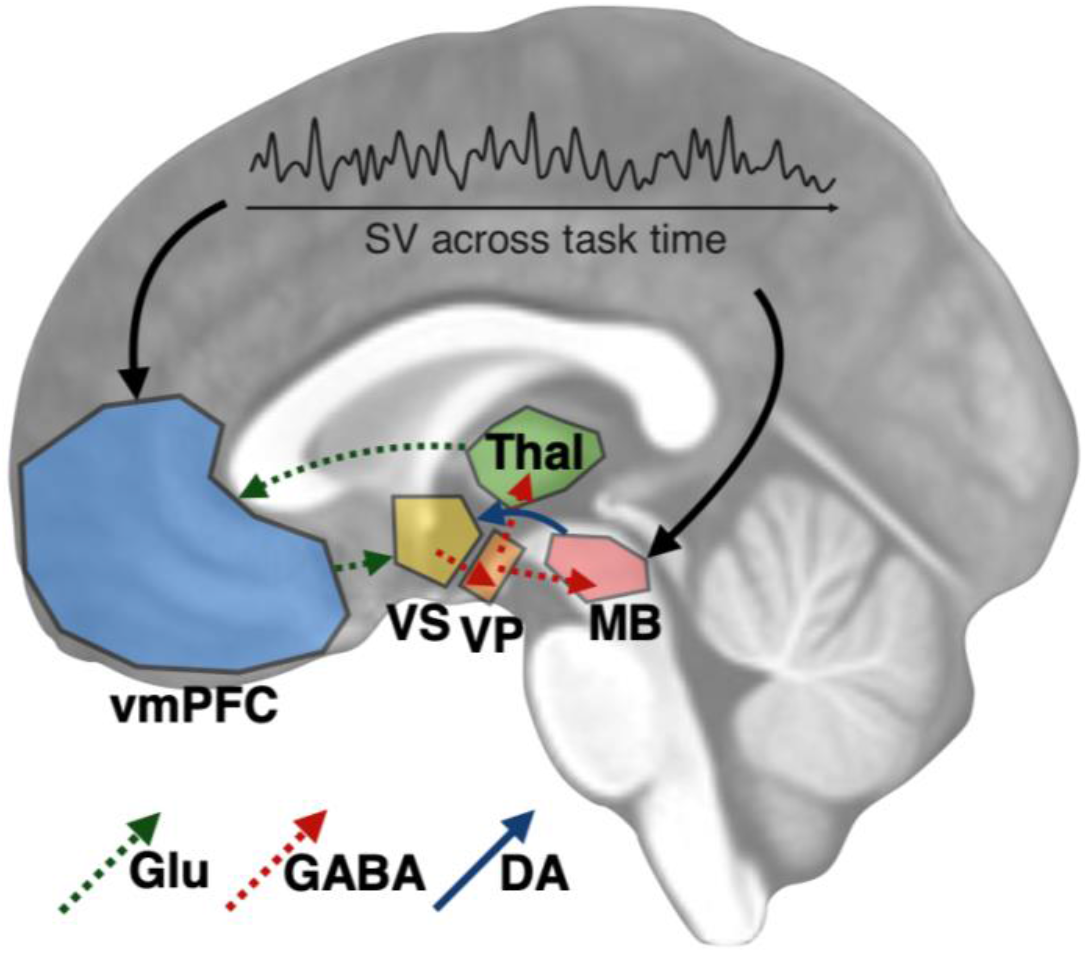
Illustration of a potential mechanism by which mesolimbic D2Rs impact subjective value (SV) and vice-versa. Binding of DA to D2Rs in the ventral striatum (VS) increases GABAergic signaling to the ventral pallidum (VP), which sends GABAergic projections to the thalamus (Thal) and midbrain (MB). GABAergic VP-MB signaling promotes DA release to the VS, while VP-Thal signaling promotes glutamate signaling in the ventromedial prefrontal cortex (vmPFC). The glutamatergic afferents from the vmPFC project to and promote local DA release in the VS.

The observed voxelwise associations between D2Rs and subjective value representations in other more lateral cortical regions are consistent with prior reported effects of dopaminergic drugs^11,50,51^. While these lateral cortical regions are not often emphasized in fMRI studies of subjective value, variability in the encoding of subjective value in the IFG and precentral gyrus has been identified in studies of effort discounting^43,52^ and risky decision making^53^. In particular, Since the IFG and precentral gyrus support inhibitory control^54^ and motor control^55^, respectively, individual differences in DA function may impact corticostriatal signaling. Specifically, increased subjective value representations in the vmPFC (mediated by VS D2R) may recruit additional resources that increase motivational vigor by facilitating direct control of goal-directed movements toward highly-valued rewards^56^.

Although this is the first study of which we are aware examining associations between PET measures of DA function, fMRI measures of value processing, and discounting behavior, a related recent study identified associations between individual differences in D2R availability in the midbrain and neural representations of expected value (i.e., reward magnitude multiplied by probability) in the ventral striatum during a simple gambling task^57^. Since expected value is an objective function and does not convey details about individual subjective utility, it is not easy to draw a straightforward comparison with the present findings. Nonetheless, subjective value signal in that study could potentially be computed after taking into account each individual’s risk preferences for a more accurate direct comparison. The other unique contribution of the present study is that we used a broader set of ROIs in a larger sample to examine potential associations across the reward circuit.

A recent study^58^ in humans identified an association between PET measures of DA D1 receptor availability in the ventral striatum and reinforcement learning-based value signals in the vmPFC (but not with reinforcement learning behavior *per se*), but the dual role of DA in learned value and motivation complicates interpretation of mesolimbic DA influences on prefrontal reward processing^59^. Specifically, it remains unclear the extent to which updated state values emerging from prediction errors across time in a learning task are similar to goal values from one-shot decisions. Nevertheless, a positive correlation between vmPFC value representations and D1Rs in the ventral striatum in that study and D2Rs in the present study suggests a more nuanced relationship between DA function and value. Although D1Rs and D2Rs have opposite effects in the direct and indirect pathways of the basal ganglia, an emerging view suggests this dichotomy is specific to the dorsal striatum and non-existent in the ventral striatum^60^. Thus, mesolimbic DA signaling in the ventral striatum could support subjective value representations effectively in the same way via D1Rs or D2Rs.

While prior studies^5^ have identified subjective value representations in the VS, we did not observe this effect in this study, consistent with a recent analysis of healthy adults using a similar task^43^. As described in a meta-analysis of different kinds of value representations^61^, it has been suggested that subjective value signals in the VS might represent reward prediction errors and not goal values (which are more strongly represented in the medial prefrontal cortex). It is possible that some delay discounting tasks might have features that increased ventral striatal sensitivity to positive prediction errors—such as larger changes in the magnitudes of presented values from trial-to-trial. Since the trial-to-trial changes in presented reward magnitudes of the present study fluctuated around a normal distribution, it is possible that we minimized ventral striatal sensitivity to strong fluctuations in values. A more liberal statistical voxel threshold did reveal average subjective value representations in the ventral striatum and caudate (see unthresholded results on NeuroVault: https://neurovault.org/collections/PDSRXDAH/).

We explored all potential associations between all PET and fMRI ROIs including the PCC due to evidence from functional neuroimaging studies for subjective value signals in the PCC^5^. We did not find significant associations between D2R in any of our ROIs and subjective value signals in the PCC. It is possible that subjective value-related neural activity observed using fMRI during discounting tasks is not as DA-mediated as in the striatum, midbrain, or vmPFC. Although the PCC is often functionally co-activated with the striatum and vmPFC, it is often not included in models of reward circuitry^14^. We also did not observe associations between vmPFC D2R and subjective value signals in any of the ROIs. It is possible that effects of prefrontal DA on discounting may be more D1R-mediated as prior work suggests D1Rs and D2Rs make dissociable contributions to specific features of discounting in rodents^62,63^.

It is striking that individual differences in DA function were correlated with neural representations of subjective value but not the behavior presumed to be influenced by regions encoding subjective value. This could suggest that DA function may impact idiosyncrasies in how choice values are computed without necessarily impacting a wide array of possible choice behaviors. The lack of a correlation between D2Rs and reward discounting is consistent across studies of healthy adults in a larger sample^2^. Together, these individual differences may suggest that revealed preferences indexed by behavioral choices are not aligned tightly enough to valuation signals indexed by BOLD responses to capture the biological mechanisms that shape valuation. Although this runs counter to assumptions in the neuroeconomics literature, the lack of strong associations complements some theoretical models of cognition. For example, fitting David Marr’s levels of analysis, DA signaling provides a neural substrate at the implementation level, subjective value provides the strategy at the algorithmic level, and preference for smaller-sooner options describe the problem at the computational level^64^. As Marr and others have described, the neural processes alone at the implementation level cannot adequately describe behavior at the computational level, but only have meaning inasmuch as each of these levels are linked by the intermediate algorithmic level^65^. This hierarchical structure might explain why D2R receptor *availability* (implementation level) alone does not reveal associations with discounting behavior (computational level), even though it does explain neural subjective value representation (algorithmic level). This precludes dynamic measures of dopamine signals (for example fast-scan cyclic voltammetry) which more directly encode value signals^9^.

As with most neuroreceptor PET studies, the most important limitation of the present study is the sample size, which limits statistical power. Although the sample size of this study is comparable to or larger than other recent studies measuring both fMRI signal and DA PET measures within subjects^57,66,67^, no prior studies have used large enough samples to better estimate the effect sizes that might be expected. As such, even the strongest effects have quite wide confidence intervals, so the sizes of the true associations between these measures are unclear. Despite the relatively small sample, given the novelty of the associations observed, these results provide valuable information given the direct measurement of DA receptors and past speculation on the role of DA in the study of reward discounting. It is also important to note that since we did not measure DA release, we are limited from making stronger claims about transient changes in VS DA concentrations. Instead, baseline measures of D2R availability reflect individual differences that are more trait-like. Nonetheless, baseline measures have previously been shown to be positively correlated with DA release^68^. Since all neuroimaging data (fMRI and PET) are publicly available on OpenNeuro (https://openneuro.org/datasets/ds002041), we hope this initial set of analyses and the complete data set provide a unique resource for other scientists to better understand associations between DA receptors and reward-related functional brain activation. Until now, it has been unclear how neural representations of subjective value arise to support a broad range of intertemporal choice behaviors in humans. The present findings suggest that variation dopamine function may account for differences between people in neural representations of subjective value.

## Acknowledgements

This research was supported by National Institute on Drug Abuse grant R21-DA033611.

